# *De novo* designed Hsp70 activator dissolves intracellular condensates

**DOI:** 10.1101/2023.09.18.558356

**Authors:** Jason Z Zhang, Nathan Greenwood, Jason Hernandez, Josh T Cuperus, Buwei Huang, Bryan D Ryder, Christine Queitsch, Jason E Gestwicki, David Baker

## Abstract

Protein quality control (PQC) is carried out in part by the chaperone Hsp70, in concert with adapters of the J-domain protein (JDP) family. The JDPs, also called Hsp40s, are thought to recruit Hsp70 into complexes with specific client proteins. However, the molecular principles regulating this process are not well understood. We describe the *de novo* design of a set of Hsp70 binding proteins that either inhibited or stimulated Hsp70’s ATPase activity; a stimulating design promoted the refolding of denatured luciferase *in vitro*, similar to native JDPs. Targeting of this design to intracellular condensates resulted in their nearly complete dissolution. The designs inform our understanding of chaperone structure-function relationships and provide a general and modular way to target PQC systems to condensates and other cellular targets.

## Introduction

Heat shock protein 70 (Hsp70) is a central component of protein quality control (PQC), playing roles in protein folding, trafficking and turnover^1^. A key step in this process is Hsp70 recruitment to diverse “client” proteins by members of the J-domain protein (JDP) family of adapters^2^. There are 45+ JDPs in humans and studies have started to reveal which ones are responsible for specific functions^3,4^. However, it has been proven challenging to ascertain the roles of many JDPs, likely due to partial functional redundancies. To unravel the complexity of this system, orthogonal approaches are needed.

Towards that goal, we considered the *de novo* design of a modular, artificial JDP. The prototypical human JDP, DnaJA2, is composed of an N-terminal J-domain (JD), a glycine/phenylalanine (G/F)-rich region, two C-terminal domains (CTDI/II) and a dimerization motif. The JD is a conserved, four-helix bundle that displays a histidine-proline-aspartate (HPD) motif that is invariant across eukaryotic and prokaryotic organisms^5^. The JD interacts with Hsp70 between the nucleotide binding domain (NBD) and the substrate-binding domain (SBD) (**Fig 1a**)^5^, using its HPD motif to accelerate Hsp70’s ATPase activity. In turn, this step triggers large conformational changes in Hsp70 and promotes its tight binding to client proteins^6^. In this way, the JDP not only recruits Hsp70 to clients, but also activates the chaperone in the vicinity of intended client proteins. Another key feature of many JDPs, such as DnaJA2, is that their CTDI/II domains interact with subsets of client proteins^7^, engaging in their “hand-off” to Hsp70. Thus, the *de novo* design of an artificial JDP poses interesting challenges because it would need to mimic: (i) binding to an evolutionarily invariant site on Hsp70, (ii) activation of a complex, ATPase driven conformational change and (iii) coordinated client “hand-off”. We anticipated that seeking to design JDPs de novo could provide molecular insights into the function of the natural PQC systems and open up new opportunities for targeting Hsp70 foldase activity to specific targets.

**Figure 1:**
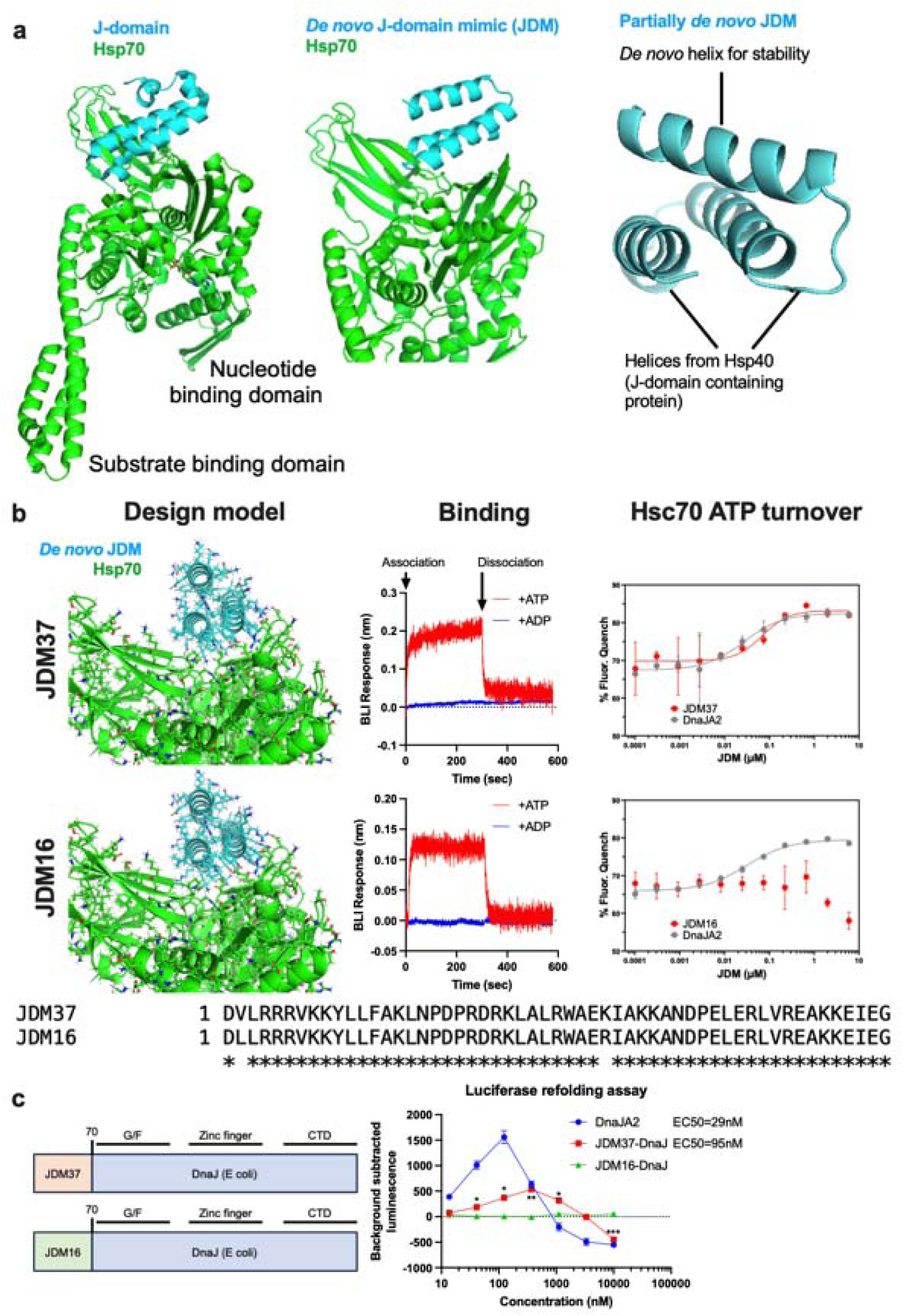
*De novo* design of Hsp70 binders/activators. **a**, Structures of Hsp70 with either native or designed proteins. Left: Crystal structure of DnaK (Hsp70) with DnaJ (Hsc40) (PDB: 5NRO). Middle: AlphaFold structure prediction of DnaK with fully *de novo* protein designed to bind to the same region as DnaJ. Right: AlphaFold structure prediction of DnaK with partially redesigned DnaJ. **b**, Design and biochemical characterization of fully *de novo* designed J-domain mimics (JDMs). Left: AlphaFold complex prediction of selected JDM with DnaK. Middle: Representative trace of biolayer interferometry (BLI) measurements of JDM binding to either ATP or ADP-loaded Hsc70. Right: *In* vitro assays measuring ATP turnover by Hsc70 with either native DnaJA2 (Hsp40) or JDMs. Lines represent average of fluorescence quenching (n=3 experiments). Bottom: Sequence alignment of the two JDMs presented here. **c**, *In vitro* assays measuring the refolding of denatured luciferase where Hsc70 and either native DnaJA2 (Hsp40) or JDM37 fused to DnaJ without its native J-domain (JDM37-DnaJ) was added. Lines represent average luminescence recorded (n=3 experiments).

## Results

### Computational design of Hsp70 binders

We set out to design *de novo* proteins which we call J-domain mimics (JDMs) that bind to Hsp70 at the same site as the native JD (**Fig. 1a**). We employed Rosetta protein design to construct JDMs in two ways (**Fig. 1a and Extended Data Fig. 1a**, see Methods for details). First^8^, we designed fully *de novo* JDMs that are predicted to bind Hsp70 at the same site as native JDs but are unrelated in structure and sequence. Second^9^, we built upon the two helices within DnaJ (a well-studied *E. coli* JDP) that interact with its partner (DnaK, the *E. coli* Hsp70) and added a third, designed helix to improve binding and monomer stability. In brief^8^, over a million designs were generated *in silico* and filtered based on their predicted binding to DnaK using Rosetta and AlphaFold2 (AF2) metrics (see Methods for details). Then, 20,000 of the fully *de novo* and 5,000 of the partially *de novo* JDM designs were displayed on the surface of yeast and sorted based on binding to biotinylated Hsc70 (human HSPA9). From this screen, 4 fully *de novo* JDM designs bound to Hsc70, while none of the partially *de novo* JDMs did (**Extended Data Fig. 1b**). These four designs were then affinity matured through site-saturation mutagenesis (SSM) (**Extended Data Fig. 1b-c**) followed by combination of the positive variants into optimized JDMs. This process resulted in 41 JDM design variants that showed strong binding to immobilized Hsc70 at low concentration (10 nM). Like native JDs, the JDMs bound only to Hsc70 when bound to ATP, but not ADP, and they could be displaced by addition of a molar excess of a native JDP (human DnaJB1) (**Extended Data Fig. 1d**), suggesting that they bind the intended site.

### Screening for active JDM designs

We purified the 41 JDMs from *E. coli* and confirmed their binding to Hsp70 by biolayer interferometry (BLI) (**Extended Data Fig. 2–4**), revealing that the designs had affinities that ranged from 1nM to 1µM. In alignment with the previous data (**Extended Fig. 1d**), the purified JDMs bound to Hsc70 in the ATP-bound state but lacked binding to the ADP-loaded Hsc70 (**Fig. 1b**). Next, we investigated whether the designs could promote steady-state ATP turnover. The 30 well expressing JDMs were tested at a range of concentrations for the ability to stimulate Hsc70 (1 µM) in ATP turnover assays (**Extended Data Fig. 5a**). While most inhibited Hsc70’s ATP turnover, one of the designs, JDM37, promoted Hsc70’s ATP hydrolysis, with EC_50_ and maximum turnover values that are comparable to the natural protein, DnaJA2 (**Fig. 1b and Extended Data Fig. 5a**). To test this design *in vivo*, we expressed JDM37 in *E. coli* and measured cell growth during 42°C heat stress compared to empty vector (EV) control. The JD-binding site on human Hsc70 and *E. coli* DnaK is highly conserved (~80% similar), so we reasoned that good designs might partially support growth under these conditions. Indeed, JDM37 increased *E. coli* growth rate at 42°C (**Extended Data Fig. 6a-b**). Conversely, *E. coli* expressing many of the other JDMs slowed growth rate during heat stress. This finding suggests that some of the designs might bind Hsp70s, but with kinetics that are not compatible with promoting ATPase activity. As predicted, the most potent suppressors of *E. coli* growth at 42°C competed with the ability of DnaJA2 to activate Hsc70’s ATPase activity *in vitro* (**Extended Data Fig. 6c**). Of note, one of these competitive inhibitors, JDM16, differs in only two amino acids from the successful JDM37 design (**Fig. 1b**). When we tested purified JDM16 for binding, it selectively interacted with ATP-loaded Hsc70, but it did not induce Hsc70’s ATP turnover (**Fig. 1b**). Overall, we have generated artificial JDs that act either as Hsp70 activators (*i.e.,* JDM37) or inhibitors (*i.e.,* JDM16).

The refolding of chemically denatured luciferase is a complex process that requires coordination of client hand-off during multiple cycles of ATPase activity^10,11^. This process is known to require not only the JD, but also the CTD1/II of natural JDPs^12^. Not surprisingly, we found that purified JDM37 alone does not promote Hsc70-mediated luciferase refolding (**Extended Data Fig. 5b**), as it lacks client binding activity. However, when we created a chimera in which the natural JD of DnaJB1 is replaced by JDM37, we found that it collaborated with Hsc70 to mediate luciferase refolding (**Fig. 1c**). Although the activity of the JDM37-chimera was ~3 fold less than that of native Hsp40 (DnaJA2) and the EC_50_ was ~3-fold higher, this result suggested that artificial JDPs can indeed engage in complex chaperone functions. Compared to DnaJA2, the JDM37 chimera had a slightly weaker EC_50_ in ATP turnover assays (see **Fig. 1b**) and altered Hsc70-binding kinetics (**Extended Data Fig. 7**), which may explain the modestly weaker refolding activity. In contrast, replacing JDM37 with the inhibitor, JDM16, led to a chimera with no activity in the luciferase refolding assay (**Fig. 1c**). Thus, chimeras of *de novo* Hsp70 activators can recapitulate complex functions of JDPs.

### Dissolving intracellular condensates by *de novo* JDMs

Condensates are membrane-less organelles that are stabilized by a network of weak, multivalent protein interactions^13^, often involving contacts between disordered regions. Hsp70 has been associated with the disassembly of condensates and ATP turnover is required for this process^14,15^. Thus, one potential use of our artificial JDM technology is to build adapters that selectively recruit Hsc70 to these unique compartments. To test this idea, we first expressed the condensate-forming protein, RIα, as an mCherry (mCh) fusion in HEK293T cells. Then, we targeted JDM37 to these RIα condensates by fusing it to an mCherry nanobody (mChnb). To allow co-localization studies, we also incorporated TSapphire into the JDM37-mChnb fusion (**Fig. 2a**). We found that expression of JDM37-mChnb-TSapphire in the HEK293T cells decreased ~4-fold the number of RIα-mCh puncta per cell (**Fig. 2b and Extended Data Fig. 8a-c**). In contrast to our controls, either fusing JDM16 to mChnb and TSapphire (JDM16-mChnb-TSapphire) or fusing only mChnb with TSapphire (mChnb-TSapphire) did not decrease the number of RIα-mCh puncta per cell (**Fig. 2b**). Thus, both Hsp70 recruitment and ATP turnover are required for RIα condensate dissolution. The disassembly of the condensate was not associated with the degradation or the loss of RIα-mCh protein (**Extended Data Fig. 8a**). This is an important observation because “stalled” Hsp70 complexes are associated with turnover of some clients^16^. Next, we explored the downstream consequences of this induced dissolution of RIα condensates. Specifically, it is known that these structures sequester cAMP and active PKA^15^, so we investigated whether expression of JDM37-mChnd-TSapphire might release them. Indeed, dissolving RIα condensates by expression of JDM37-mChnb-TSapphire increased global cAMP accumulation and PKA activation after physiological isoproterenol stimulation, as evidenced by time-course ICUE3 and AKAR4 biosensor imaging (**Fig. 2c-d and Extended Data Fig. 8d-e**), respectively.

**Figure 2:**
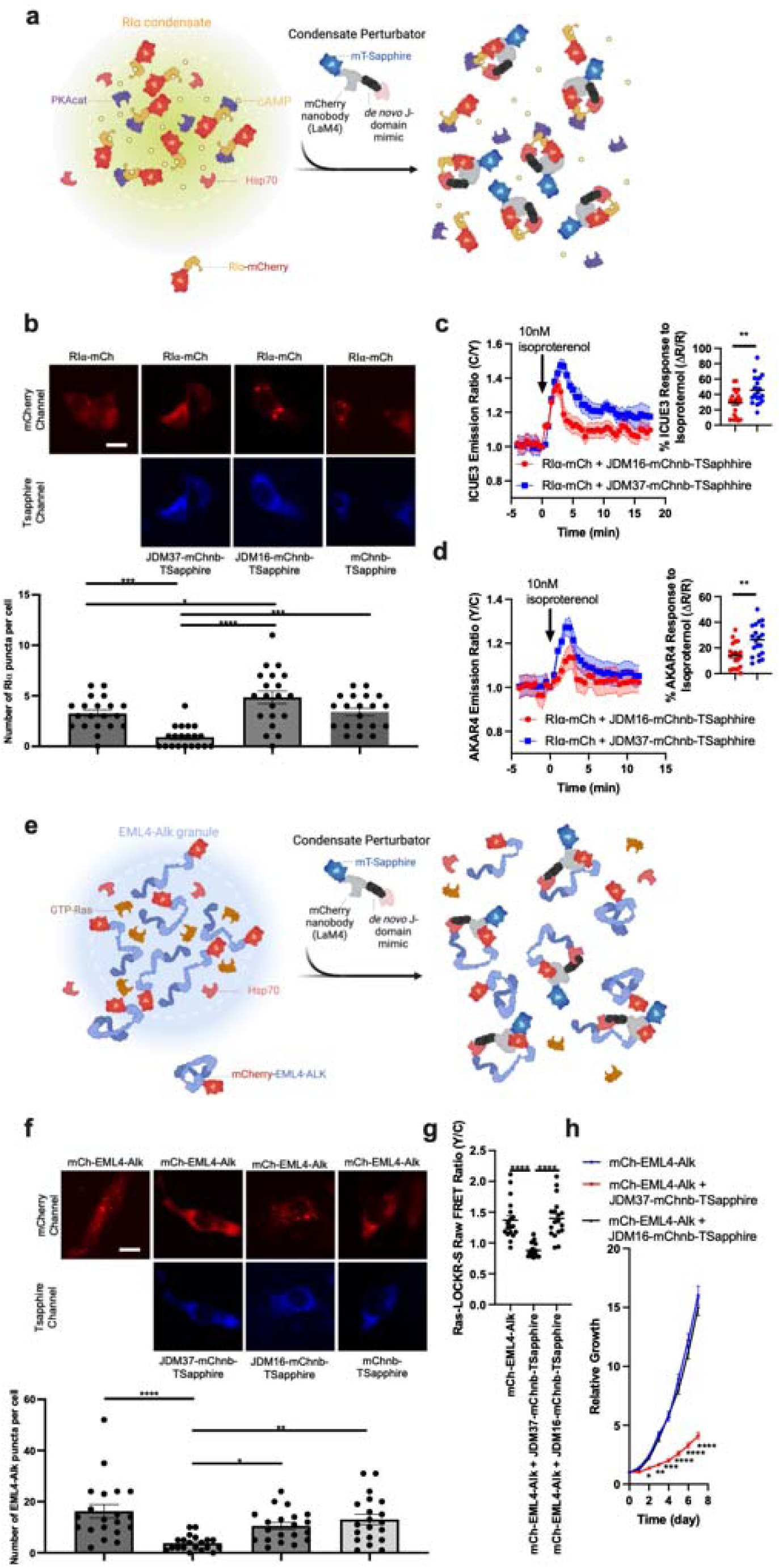
Fusing *de novo* designed JDMs with substrate binding domains dissolves intracellular condensates. **a**, Schematic of strategy to dissolve RIα condensates. Protein that drives condensation (here, RIα) is tethered with mCherry. JDM37 is fused to an mCherry nanobody and mT-Sapphire, and this tool is called condensate perturbator. **b**, Expression of engineered condensate perturbators dissolves mCherry-tagged RIα puncta in HEK293T cells (see Methods for details). Top: Representative epifluorescence images of the various conditions tested. Scale bar = 10µm. Bottom: Quantification of number of RIα puncta per cell. Each point represents a single cell (n=20 cells). **c-d**, Time-course imaging of HEK293T cells expressing mCherry-tagged RIα, either cAMP sensor ICUE3 (**c**) or PKA sensor AKAR4 (**d**), and either JDM37-based or JDM16-based condensate perturbators. In each condition, 10nM isoproterenol was added. Solid lines indicate representative average time with error bars representing standard error mean (SEM) (n=at least 15 cells per curve). **e**, Schematic of strategy to dissolve EML4-Alk oncogenic condensates. Protein that drives condensation (here, EML4-Alk) is tethered with mCherry. **f**, Expression of engineered condensate perturbators dissolves mCherry-tagged EML4-Alk puncta in Beas2B cells (see Methods for details). Top: Representative epifluorescence images of the various conditions tested. Scale bar = 10µm. Bottom: Quantification of number of EML4-Alk puncta per cell. Each point represents a single cell (n=20 cells). **g**, Raw FRET ratios of HEK293T cells expressing mCherry-tagged RIα, Ras sensor Ras-LOCKR-S, and either JDM37-based or JDM16-based condensate perturbators. Each point represents a single cell (n=18 cells). **h**, Cell growth curves of Beas2B cells expressing mCherry-tagged EML4-Alk with or without condensate perturbators (n=3 experiments). Line represents average from all 3 experiments. For the quantification of number of puncta per cell in **b and f**, cells only with sufficient expression of the condensate perturbator were chosen for analysis (see Methods for details).

The fusion oncoprotein EML4-Alk (associated with lung adenocarcinoma) can also form condensates that regulate signaling^17^. However, it is not clear whether the formation of this compartment is strictly necessary for oncogenesis. We targeted the JDM37-mChnb-TSapphire protein to mCh-tagged EML4-Alk (**Fig. 2e**) in Beas2B lung cells and observed a drastic decrease in the number of mCh-EML4-Alk puncta per cell, (**Fig. 2f and Extended Data Fig. 8f-h**). Once again, the key controls, JDM16-mChnb-TSapphire and mChnb-TSapphire, had no effect. One of the mechanisms by which EML-Alk condensates are thought to act is by locally activating Ras^17^. Consistent with this idea, dissolution of EML4-Alk puncta by JDM37-mChnb-TSapphire, but not JDM16-mChnb-TSapphire, in Beas2B cells decreased Ras signaling, as measured by a Ras-LOCKR-S biosensor^18^ (**Fig. 2g**). Expression of JDM37-mChnb-TSapphire in EML4-Alk-expressing Beas2B cells also diminished cell growth by ~4-fold compared to controls (**Fig. 2h**). These results suggest that the formation of EML4-Alk condensates is critical for oncogenic signaling and cell growth, further highlighting the suitability of this condensate as a putative drug target.

## Discussion

The Hsp70-JDP system is an ancient mediator of PQC^1^. To better understand the function of this important complex, we have designed a modular, artificial JDP. Although our major goal here was to develop the technology platform, the results have already begun to illuminate structure-function relationships. For example, JDM37 and JDM16 differ at only two positions, but only JDM37 stimulates ATPase activity (see **Fig. 1b**). The designed JDMs have discrete biochemical (**Extended Data Fig. 4**) and structural features (**Extended Data Fig. 9 and Supplementary Note**), which may bring insight into what governs the regulation of Hsp70s. Another key insight comes from the observation that chimeras of JDM37, but not JDM16, dissolved two different condensates, consistent with both recruitment and ATPase activity^19^ being required to regulate intracellular protein clients. We anticipate that the ability to target Hsp70 to specific subcellular locations and targets with substrate-specific synthetic JDPs will be broadly useful given the modularity and tunability of the system.

## Methods

### Designing partially *de novo* J-domain mimics

The crystal structure (PDB: 5NRO) of the J-domain-activated conformation of DnaK (ATP-loaded DnaK bound to the J-domain of DnaJ) was refined with the Rosetta FastRelax protocol with coordinate constraints. Miniprotein binder generation began by paring down J-domain by removing its first and fourth helix as they do not directly interact with DnaK. Afterwards, a third helix on top of the remaining helices were added by using the RosettaRemodel blueprint^9^ program in order to increase stability of the remaining helices. 30,000 blueprints that differed in the lengths of the helices and the loop types were generated. The whole miniprotein except the key interacting residues on the J-domain were then designed using the FastDesign protocol. From these designs, the monomeric metrics and the interface metrics were calculated using Rosetta and these metrics were used to filter designs based on maintaining the shape of the interacting helices (particularly the HPD motif geometry). Top 15,000 designs were selected for AlphaFold2 structure predictions in complex with DnaK and these predicted complexes were again calculated for interface metrics using Rosetta. These metrics were used to identify the top 5,000 designs based on its predicted binding to DnaK. This protocol is graphically summarized in **Extended Data Fig. 1a**.

### Designing fully *de novo* J-domain mimics

This protocol has been described previously^8^. In brief, the crystal structure (PDB: 5NRO) of the J-domain-activated conformation of DnaK (ATP-loaded DnaK bound to the J-domain of DnaJ) was refined with the Rosetta FastRelax protocol with coordinate constraints, and the J-domain was extracted for subsequent steps. Initial docking conformations were generated by RifDock. Next, billions of individual disembodied amino acids were docked against DnaK using RifDock and specifying the dock with residues (6-10) on DnaK that interact with DnaJ based on 5NRO. The ones that passed a specific energy cutoff value (−1.5 Rosetta energy unit) were stored and the corresponding inverse rotamers were generated. The de novo scaffold library of 19,000 miniproteins (in length 56 - 65 residues, mostly three helical bundles) were docked into the field of the inverse rotamers to produce initial docked conformations. These docked conformations were further optimized using the FastDesign protocol to generate shape and chemically complementary interfaces. Computational metrics of the final design models were calculated using Rosetta, which includes ddg, shape complementary, contact patch, and interface buried solvent accessible surface area. The designs that scored well in these interface metrics were analyzed for common motifs (couple amino acids in length) and these privileged motifs were fed into MotifGraft to build the next set of miniproteins. Afterwards, these next set of designs were again ran through FastDesign. 2,000,000 of these designs were then predicted in AlphaFold2 structure predictions in complex with DnaK and these predicted complexes were again calculated for interface metrics using Rosetta. Several iterations of FastDesign, MotifGraft, and AlphaFold2 were performed, as diagrammed in **Extended Data Fig. 1a**. After final AlphaFold2 predictions and metric scoring, the top 25,000 designs based on its predicted binding to DnaK were ordered.

### DNA library preparation

All protein sequences were padded to a uniform length (65 amino acids) by adding a (GGGS)n linker at the C terminal of the designs, to avoid the biased amplification of short DNA fragments during PCR reactions. The protein sequences were reversed translated and optimized using DNAworks2.0 with the S. cerevisiae codon frequency table. Homologous to the pETCON plasmid Oligo libraries encoding the designs were ordered from Twist Bioscience. Oligo pool encoding the *de novo* designs and the point mutant library for SSM were ordered from Agilent Technologies. Combinatorial libraries were ordered as IDT (Integrated DNA Technologies) ultramers with the final DNA diversity ranging from 1×10^6^ to 1×10^7^.

All libraries were amplified using Kapa HiFi Polymerase (Kapa Biosystems) with a qPCR machine (BioRAD CFX96). In detail, the libraries were firstly amplified in a 25 μL reaction, and PCR reaction was terminated when the reaction reached half the maximum yield to avoid over-amplification. The PCR product was loaded to a DNA agarose gel. The band with the expected size was cut out and DNA fragments were extracted using QIAquick kits (Qiagen, Inc.). Then, the DNA product was re-amplified as before to generate enough DNA for yeast transformation. The final PCR product was cleaned up with a QIAquick Clean up kit (Qiagen, Inc.). For the yeast transformation, 2-3 μg of digested modified pETcon vector (pETcon3) and 6 μg of insert were transformed into EBY100 yeast strain using the protocol as described before.

DNA libraries for deep sequencing were prepared using the same PCR protocol, except the first step started from yeast plasmid prepared from 5×10^7^ to 1×10^8^ cells by Zymoprep (Zymo Research). Illumina adapters and 6-bp pool-specific barcodes were added in the second qPCR step. Gel extraction was used to get the final DNA product for sequencing. All libraries include the native library and different sorting pools were sequenced using Illumina NextSeq/MiSeq sequencing.

All pcDNA3.1 plasmids constructed here were produced by GenScript.

### Yeast surface display

*S. cerevisiae* EBY100 strain cultures were grown in C-Trp-Ura media and induced in SGCAA media following the protocol as described before. Cells were washed with PBSF (PBS with 1% BSA) and labelled with biotinylated Hsc70 using two labeling methods, with-avidity and without-avidity labeling. For the with-avidity method, the cells were incubated with biotinylated Hsc70, together with anti-c-Myc fluorescein isothiocyanate (FITC, Miltenyi Biotech) and streptavidin–phycoerythrin (SAPE, ThermoFisher). The concentration of SAPE in the with-avidity method was used at 1/4 concentration of the biotinylated Hsc70. The with-avidity method was used in the first few rounds of screening of the original design to fish out weak binder candidates. For the without-avidity method, the cells were firstly incubated with biotinylated Hsc70, washed, secondarily labeled with SAPE and FITC. For these designs, three rounds of with-avidity sorts were applied at 1 μM concentration of Hsc70, 5 mM ATP, 1 μM NR peptide (NRLLLTG^20^, with N-terminal acetylation) to occupy Hsc70’s substrate binding domain, 5 mM MgCl_2_, and 10 mM KCl. For the original library of *de novo* designs, the library was sorted twice using the with-avidity method at 1 μM Hsc70, followed by several without-avidity sorts in the third round of sorting with Hsc70 concentrations at 1 μM, 100 nM and 10 nM. The SSM library was screened using the without-avidity method for four rounds, with Hsc70 concentrations at 1 μM, 100 nM, 10 nM and 1 nM. The combinatorial libraries were sorted to convergence by decreasing the target concentration with each subsequent sort and collecting only the top 0.1% of the binding population. The final sorting pools of the combinatorial libraries were sequenced using Illumina NextSeq/MiSeq sequencing. All FACS data was analyzed in FlowJo.

### Biolayer interferometry

Biolayer interferometry binding data were collected in an Octet RED96 (ForteBio) and processed using the instrument’s integrated software. For minibinder binding assays, biotinylated Hsc70 was loaded onto streptavidin-coated biosensors (SA ForteBio) at 50 nM in binding buffer (10 mM HEPES (pH 7.4), 150 mM NaCl, 3 mM EDTA, 0.05% surfactant P20, 0.5% non-fat dry milk) along with 5 mM ATP, 1 μM NR peptide, 5mM MgCl_2_, and 10 mM KCl for 360 s. Analyte proteins were diluted from concentrated stocks into binding buffer. After baseline measurement in the binding buffer alone, the binding kinetics were monitored by dipping the biosensors in wells containing the target protein at the indicated concentration (association step) and then dipping the sensors back into baseline/buffer (dissociation). Data were analyzed and processed using ForteBio Data Analysis software v.9.0.0.14.

### Cell culture and transfection

HEK293T cells were cultured in Dulbecco’s modified Eagle medium (DMEM) containing 1 g L^-1^ glucose and supplemented with 10% (v/v) fetal bovine serum (FBS) and 1% (v/v) penicillin–streptomycin (Pen-Strep). Beas2B cells were cultured in Roswell Park Memorial Institute 1640 (RPMI 1640) with 10% (v/v) FBS and 1% Pen-Strep. All cells were grown in a humidified incubator at 5% CO_2_ and at 37°C.

Before transfection, all cells were plated onto sterile poly-D-lysine coated plates or dishes and grown to 50%–70% confluence. HEK293T and Beas2B cells were transfected using Turbofectin 8 and grown for an either an additional 16-24 hours for HEK293T or 48 hours for Beas2B before imaging. Beas2B cells underwent serum starvation for 16-24 hours prior to downstream experiments.

### General procedures for bacterial protein production and purification

For all the JDMs, DnaJB1, and Hsc70, the *E. coli* Lemo21(DE3) strain was transformed with a pET29b^+^ plasmid encoding the synthesized gene of interest. Cells were grown for 24 hour in liquid broth medium supplemented with kanamycin. Cells were inoculated at a 1:50 mL ratio in the Studier TBM-5052 autoinduction medium supplemented with kanamycin, grown at 37°C for 2–4 hours and then grown at 18°C for an additional 18 hours. Cells were collected by centrifugation at 4,000 *g* at 4°C for 15 min and resuspended in 30 mL lysis buffer (20 mM Tris-HCl, pH 8.0, 300 mM NaCl, 30 mM imidazole, 1 mM PMSF and 0.02 mg ml^-1^ DNase). Cell resuspensions were lysed by sonication for 2.5 min (5 s cycles). Lysates were clarified by centrifugation at 24,000 xg at 4°C for 20 min and passed through Ni-NTA nickel resin (2 mL) pre-equilibrated with wash buffer (20 mM Tris-HCl, pH 8.0, 300 mM NaCl and 30 mM imidazole). The resin was washed twice with 10 column volumes of wash buffer, and then eluted with elution buffer (20 mM Tris-HCl, pH 8.0, 300 mM NaCl and 300 mM imidazole). The eluted proteins were concentrated using Ultra-15 Centrifugal Filter Units and further purified by using either Superdex 75 Increase 10/300 GL or Superdex 200 Increase 10/300 GL (depending on kDa of protein) size exclusion column in TBS (25 mM Tris-HCl, pH 8.0, and 150 mM NaCl). Fractions containing monomeric protein were pooled, concentrated and snap-frozen in liquid nitrogen and stored at −80°C.

Biotinylation of purified Hsc70 was performed using the BirA bulk kit according to manufacturer’s protocol (Avidity LLC). Briefly, biotinylation reactions (pH 8.0; 1:1 ratio) were performed overnight at 4°C on an orbital shaker and then excess biotinylation reagent was removed using Superdex 200 Increase 10/300 GL (depending on kDa of protein) size exclusion column in TBS (25 mM Tris-HCl, pH 8.0, and 150 mM NaCl).

To purify the JDMs fused to DnaJ, the same protocol as above was performed, except the cells were inoculated in Terrific Broth II buffer (TBII powder, MP Biomedicals) supplemented with kanamycin, grown at 37°C until OD_600_ 0.5 to 1, IPTG (0.2 mM) was added, and then the bacteria were grown at 37°C overnight. Cells were also lysed in BugBuster Protein Extraction buffer (EMD Millipore) for 30 min instead of sonication.

### Hsc70 ATP turnover assay

Hsc70’s ATPase activity was measured via a quinaldine red-based fluorescence assay as described before^21^. In brief, a dilution series of either DnaJA2 or JDMs was prepared and added to Hsc70 (1 µM) in 384-well plates using a Multidrop dispenser (Thermo Fisher Scientific, Inc.). Quinaldine red solution was freshly prepared each day as a 2:1:1:2 ratio of 0.05% w/v quinaldine red, 2% w/v polyvinyl alcohol, 6% w/v ammonium molybdate tetrahydrate in 6 M HCl. To each well of a 384-well white, low-volume, polystyrene plate (Greiner Bio-One, Monroe, NC) was added 5 µL of the chaperone mixture. The assay buffer (100 mM Tris, 20 mM KCl, 6 mM MgCl_2_, pH 7.4), was supplemented with 0.01% Triton X-100 to avoid identifying aggregation. Then, 5 µL of 5 mM ATP was added to initiate the reaction. The plates were centrifuged briefly, and subsequently incubated at 37 °C for 1 to 2 hrs. After incubation, 15 µL of quinaldine red solution was added, and after a 15 minute incubation, 2 µL sodium citrate (32% w/v) was added to quench the reaction. After another 15-min incubation period at 37 °C, fluorescence (excitation 430 nm, emission 530 nm) was measured in a PHERAstar plate reader.

### Luciferase refolding assay

Luciferase refolding assays followed a previously described procedure^21^. In brief, native firefly luciferase (Promega) was denatured in 8 M guanidine hydrochloride for at least 1 hour at room temperature and then diluted into assay buffer (28 mM HEPES, pH 7.6, 120 mM potassium acetate, 12 mM magnesium acetate, 2.2 mM dithiothreitol, 8.8 mM creatine phosphate, and 35 units/ml creatine kinase). Solutions were prepared of test samples (e.g. DnaJA2, JDMs), Hsc70, denatured luciferase (at 0.1 µM), and 1 mM ATP. Total volume was 25 µL in white 96-well plates (Corning Glass), and incubation time was 1 hour at 37 °C. Steady Glo reagent was prepared fresh and added to the plate immediately prior to reading luminescence.

### Bacterial cell growth assays

Single colonies from freshly transformed cells were used to inoculate 5 mL cultures of LB media containing the antibiotics required to maintain plasmids. For testing effects of JDM on WT *E. coli*, the *E. coli* Lemo21(DE3) strain was transformed with a pET29b^+^ plasmid encoding the synthesized gene of interest and cells were then grown for 24 hour in liquid broth medium supplemented with kanamycin. For growth assays, 4 μL of resuspended cells (~1 × 106 cells) were used to inoculate each well of a 96-well plate containing 196 μL of LB media (supplemented with ampicillin, and 200uM IPTG) The plate was covered with transparent LightCycler480 Sealing Foil (Roche), and grown for 24 h in a Biotek Synergy H1 plate reader at 42 °C with double orbital shaking, measuring OD_600_ every 10 min. Max-V was determined by measuring maximal growth over 15 measurements.

### Cell counting to measure cell proliferation

Beas2B cell lines were seeded in 6-wells plates at 10,000 cells/well. Cell numbers were quantified using a hemacytometer each day for 7 days.

### Epifluorescence imaging

Cells were washed twice with FluoroBrite DMEM imaging media and subsequently imaged in the same media in the dark at room temperature. Isoproterenol was added as indicated. Epifluorescence imaging was performed on a Yokogawa CSU-X1 spinning dish confocal microscope with either a Lumencor Celesta light engine with 7 laser lines (408, 445, 473, 518, 545, 635, 750 nm) or a Nikon LUN-F XL laser launch with 4 solid state lasers (405, 488, 561, 640 nm), 40x/0.95 NA objective and a Hamamatsu ORCA-Fusion scientific CMOS camera, both controlled by NIS Elements 5.30 software (Nikon). The following excitation/FRET filter combinations (center/bandwidth in nm) were used: CFP: EX445 EM483/32, CFP/YFP FRET: EX445 EM542/27, YFP: EX473 EM544/24, GFP: EX473 EM525/36, RFP (mCherry): EX545 EM605/52, TSapphire: EX405 EM525/36. All epifluorescence experiments were subsequently analyzed using Image J.

For end-point cell imaging, exposure times were 100ms for each channel with no EM gain set and no ND filter added. For time-lapse biosensor imaging, exposure times were 100 ms for acceptor direct channel (YFP) and 500ms for all other channels (CFP/YFP FRET and CFP), with no EM gain set and no ND filter added. Cells that were too bright (acceptor channel intensity is 3 standard deviations above mean intensity across experiments) or with significant photobleaching prior to drug addition were excluded from analysis.

### FRET biosensor analysis

Raw fluorescence images were corrected by subtracting the background fluorescence intensity of a cell-free region from the FRET intensities of biosensor-expressing cells. Cyan/yellow FRET ratios were then calculated at each time point (*R)*. For some curves, the resulting time courses were normalized by dividing the FRET ratio at each time point by the basal ratio value at time zero (*R*/*R*_0_), which was defined as the FRET ratio at the time point immediately preceding drug addition (*R*_0_). Graphs were plotted using GraphPad Prism 8 (GraphPad).

### Quantification of cellular puncta

For analysis of puncta number, cell images were individually thresholded and underwent particle analysis with circularity and size cutoffs in ImageJ. For the quantification of number of puncta per cell in **Fig. 2b and f**, cells only with sufficient expression of the condensate perturbator were chosen for analysis. In practical terms, with our set up, only cells with TSapphire fluorescence intensity of 1000 (arbitrary units) were included in our analysis.

### Statistics and reproducibility

No statistical methods were used to predetermine the sample size. No sample was excluded from data analysis, and no blinding was used. All data were assessed for normality. For normally distributed data, pairwise comparisons were performed using unpaired two-tailed Student’s t tests, with Welch’s correction for unequal variances used as indicated. Comparisons between three or more groups were performed using ordinary one-way or two-way analysis of variance (ANOVA) as indicated. For data that were not normally distributed, pairwise comparisons were performed using the Mann-Whitney U test, and comparisons between multiple groups were performed using the Kruskal-Wallis test. All data shown are reported as mean ± SEM and error bars in figures represent SEM of biological triplicates unless indicated otherwise. All data were analyzed and plotted using GraphPad Prism 8 including non-linear regression fitting.

## Data availability

The data that support the findings of this study are available from the corresponding author upon reasonable request. All accession codes have been provided for the paper. Source data are provided with this paper.

## Code availability

Source data are provided with this paper. Code used in this paper are provided in previous work.

## Acknowledgements

We acknowledge funding from HHMI (J.Z.Z. and D.B.), Helen Hay Whitney Foundation (J.Z.Z.), the Audacious Project at the Institute for Protein Design (J.Z.Z, D.B.) and the NIH (NS059690 to J.E.G.). We thank X. Wang, A. Motmaen, and N. Bennett for their help in computational design, S.K. Williams for providing biotinylated Hsc70, I.C. Haydon for providing some schematics in the manuscript, and M.M. Schneider for providing useful feedback on the project.

## Author contributions

J.Z.Z. conceived of the project. J.Z.Z., J.E.G., and D.B. supervised, designed, and interpreted the experiments. J.Z.Z. designed the binders with the help from B.H. for the partially *de novo* JDMs. J.Z.Z. and N.G. purified and tested the binders. J.H. tested binders in the *in vitro* activity assays. J.Z.Z. performed the mammalian cell experiments. J.T.C. and C.Q. designed and performed the *E. coli* growth experiments. B.D.R. helped with discussion for the sequence and structure comparisons. J.Z.Z., J.E.G, and D.B. wrote the original draft. All authors reviewed and commented on the manuscript.

## Competing interest

The authors claim no competing interests.

## Figure and Figure Legends

**Extended Data Figure 1:**
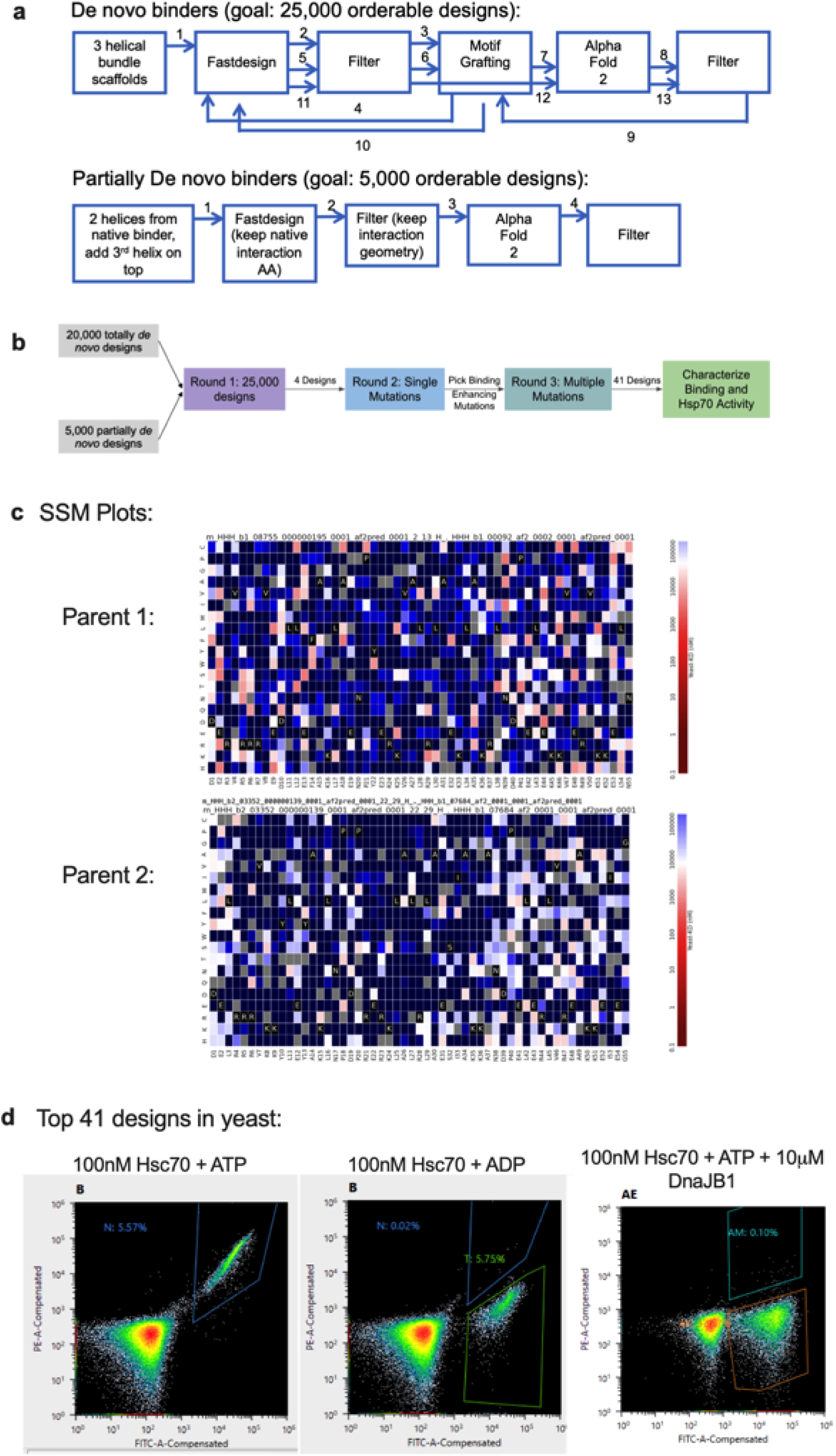
Computational and experimental pipeline for designing JDMs. **a**, Schematic of the exact computational workflow for designing fully *de novo* and partially *de novo* JDMs. **b**, Summary of the experimental (yeast display) pipeline for high throughput testing computationally designed JDMs. Round 1: computer generated designs were tested through three rounds of yeast display sorting (see Methods). Round 2: The 4 fully *de novo* JDMs were enriched and each of these JDMs underwent an SSM with three rounds of yeast display sorting. Round 3: Beneficial mutations identified through SSM were incorporated into combinatorial mutant libraries and underwent three rounds of yeast display sorting. At the end, the collected yeast cells were sequenced and the top 41 JDM designs were expressed, purified, and tested biochemically. **c**, The top 41 JDM designs come from two parents originally designed in Round 1. Shown are the two parent’s SSM maps. Each cell is a particular point mutant. Red indicates tight affinity to Hsc70 and blue indicates weak affinity to Hsc70. **d**, The top 41 JDM designs were transformed into yeast and were tested for specificity to ATP-loaded state of Hsc70 and binding to same site as native J-domains through DnaJB1 competition assays. Shown are the flow cytometry plots.

**Extended Data Figure 2:**
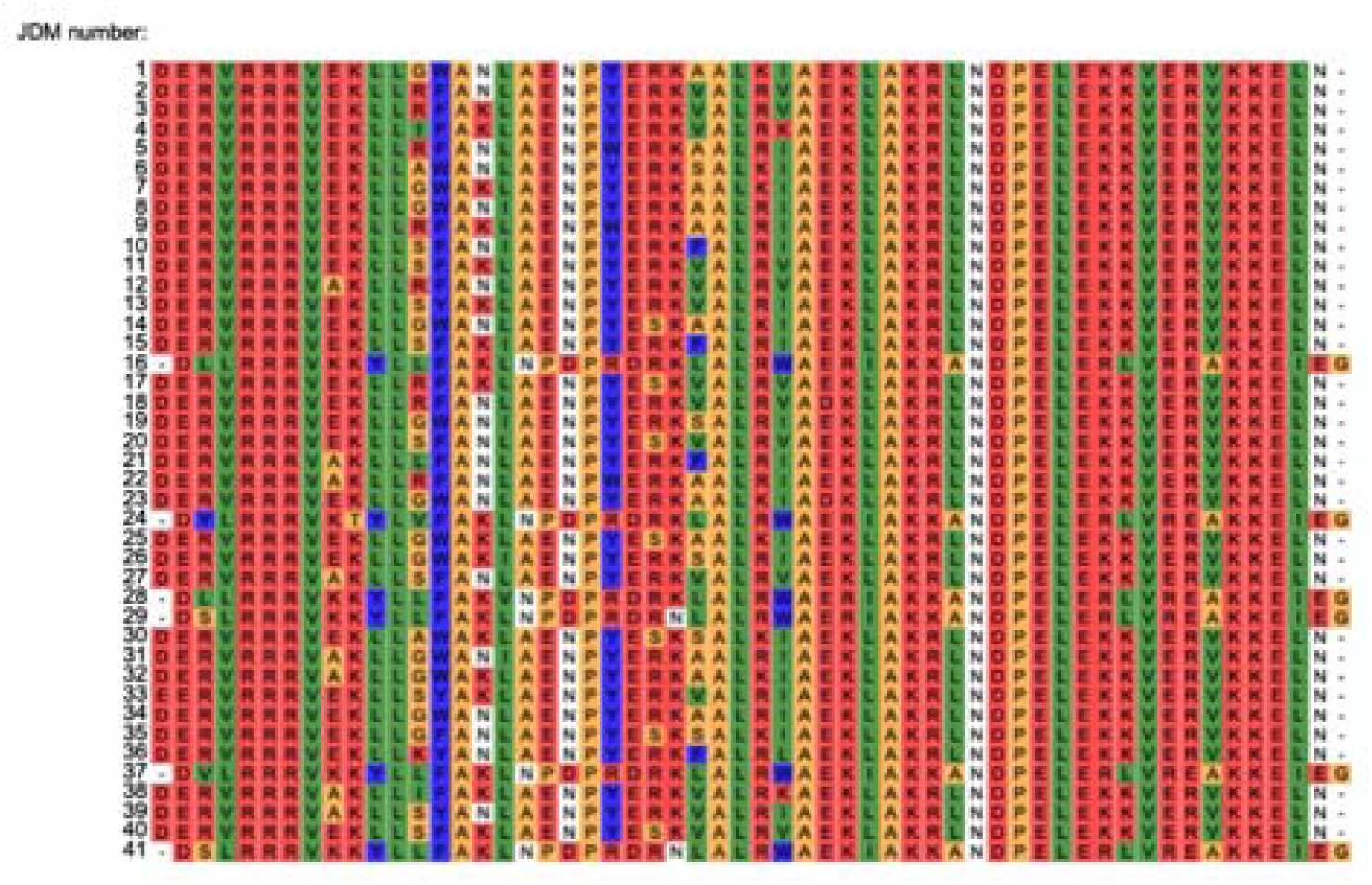
Sequences of top 41 JDM designs. Sequence alignment of the top 41 JDM designs after yeast display.

**Extended Data Figure 3:**
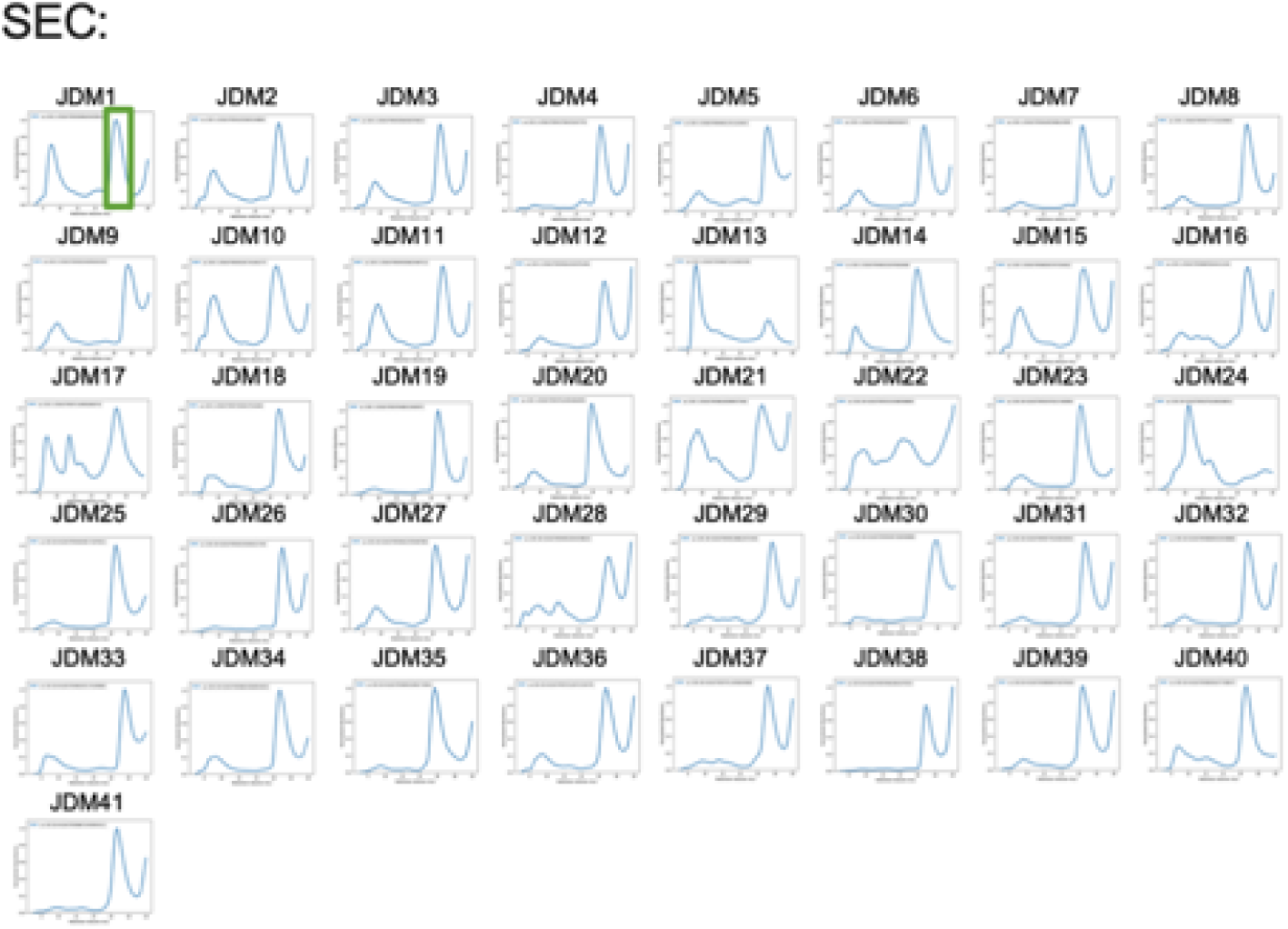
Production and purification of top 41 JDM designs. The top 41 JDM designs were transformed into *E. coli.*, induced for expression, and purified. SEC traces of the top 41 JDM designs for further purification and characterization for their size and monodispersity.

**Extended Data Figure 4:**
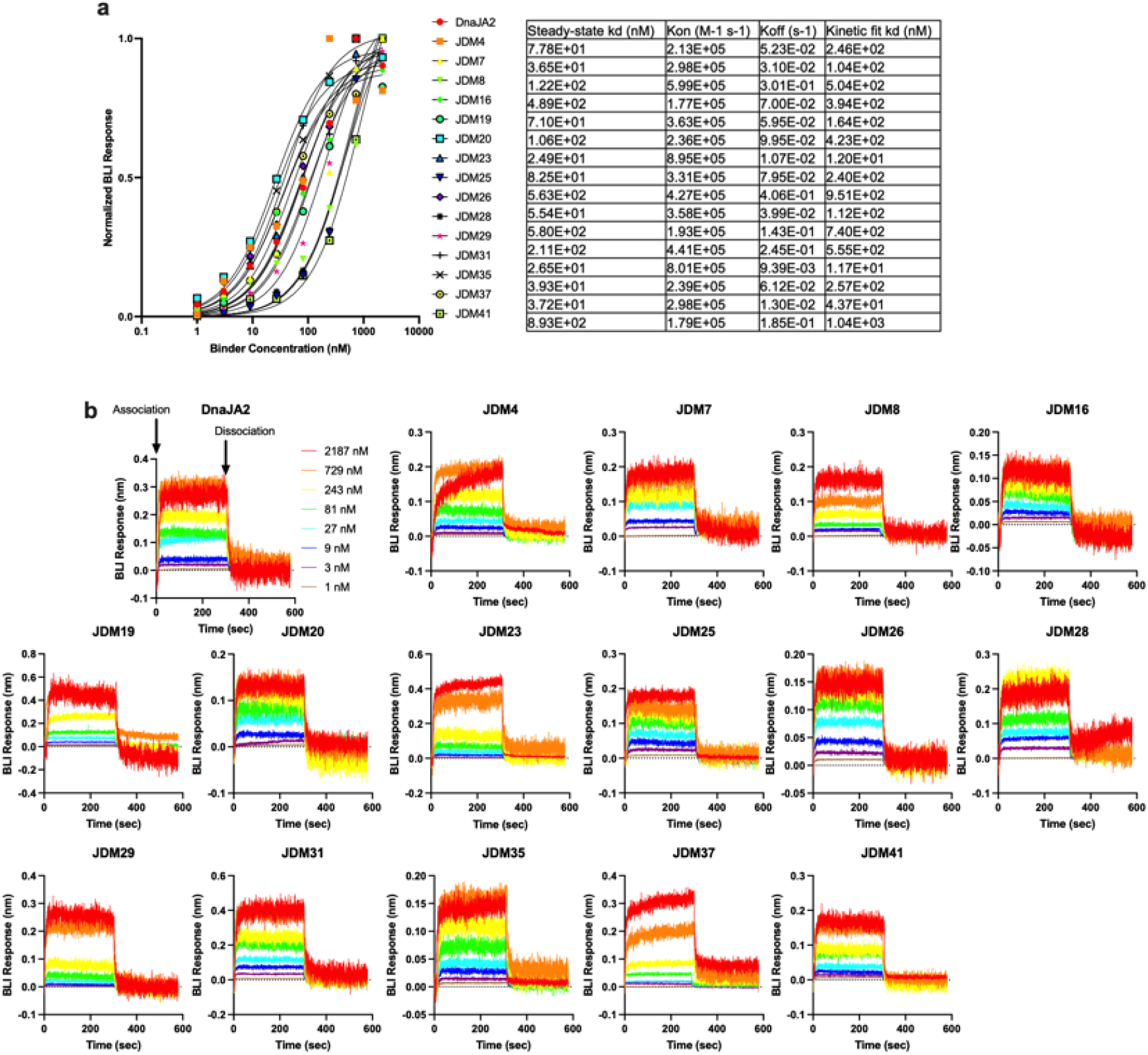
Measuring the binding of JDMs to Hsc70. **a-b**, A subset of the top 41 JDM designs (high protein yield and monodispersity) were tested in Hsc70 ATP turnover were also tested for their binding to ATP-loaded Hsc70 via BLI. Shown is the average steady-state binding titration of all the binders tested (n=3 experiments, line is EC50 curve fitting) (**a**) or a representative raw kinetic binding data (**b**).

**Extended Data Figure 5:**
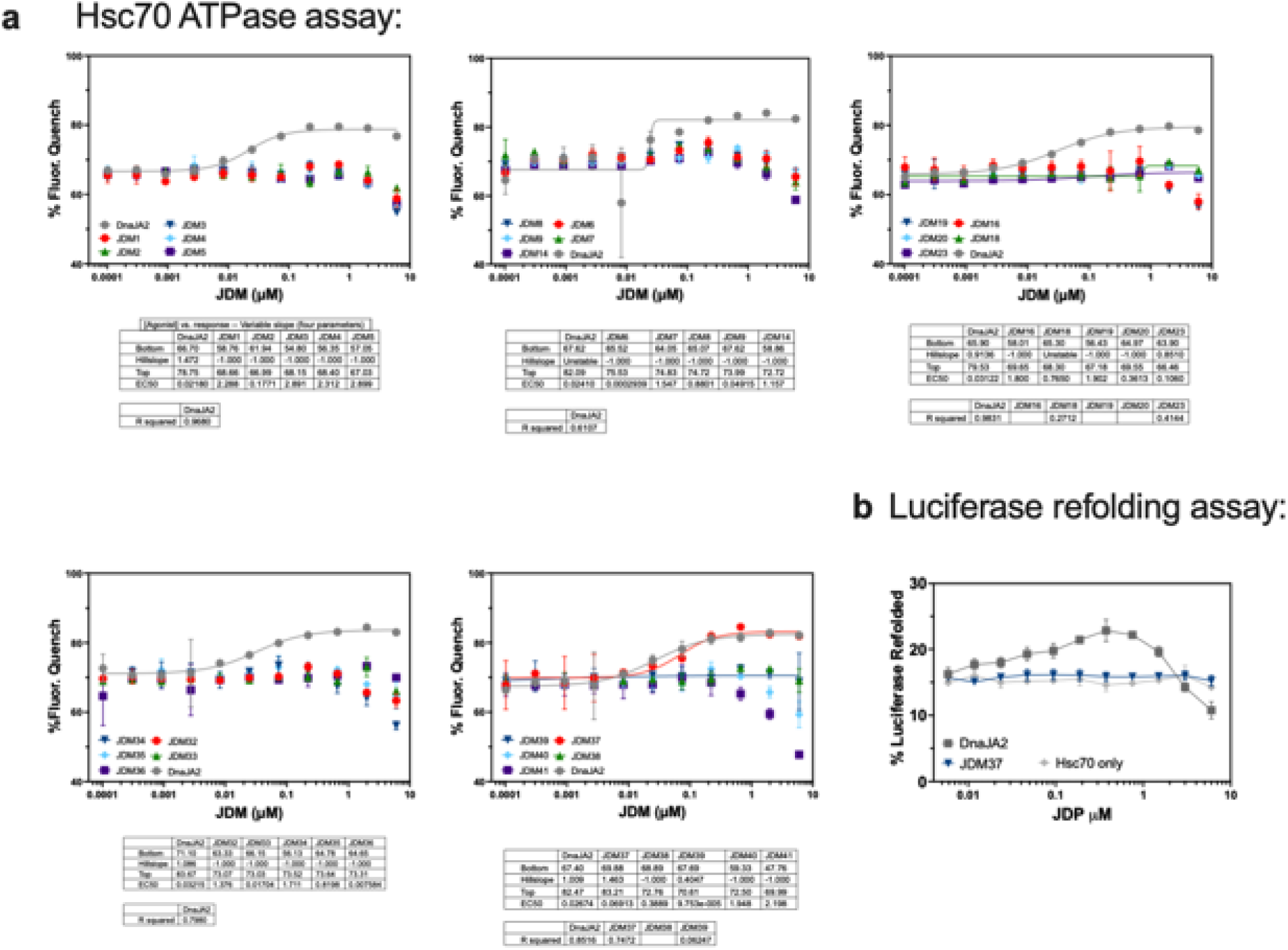
Testing JDMs effect on Hsc70 ATP turnover and refolding. **a**, A subset of the top 41 JDM designs with sufficient protein after purification and monodisperse SEC traces were tested in Hsc70 ATP turnover assays. Native DnaJA2 (Hsp40) was tested as a positive control. Lines represent average of fluorescence quenching (n=2 experiments for JDM1-5, the rest n=3 experiments). **b**, JDM37, the only JDM that activated Hsc70 ATP turnover, was tested for luciferase refolding. Native DnaJA2 (Hsp40) was tested as a positive control. Lines represent average luminescence recorded (n=3 experiments).

**Extended Data Figure 6:**
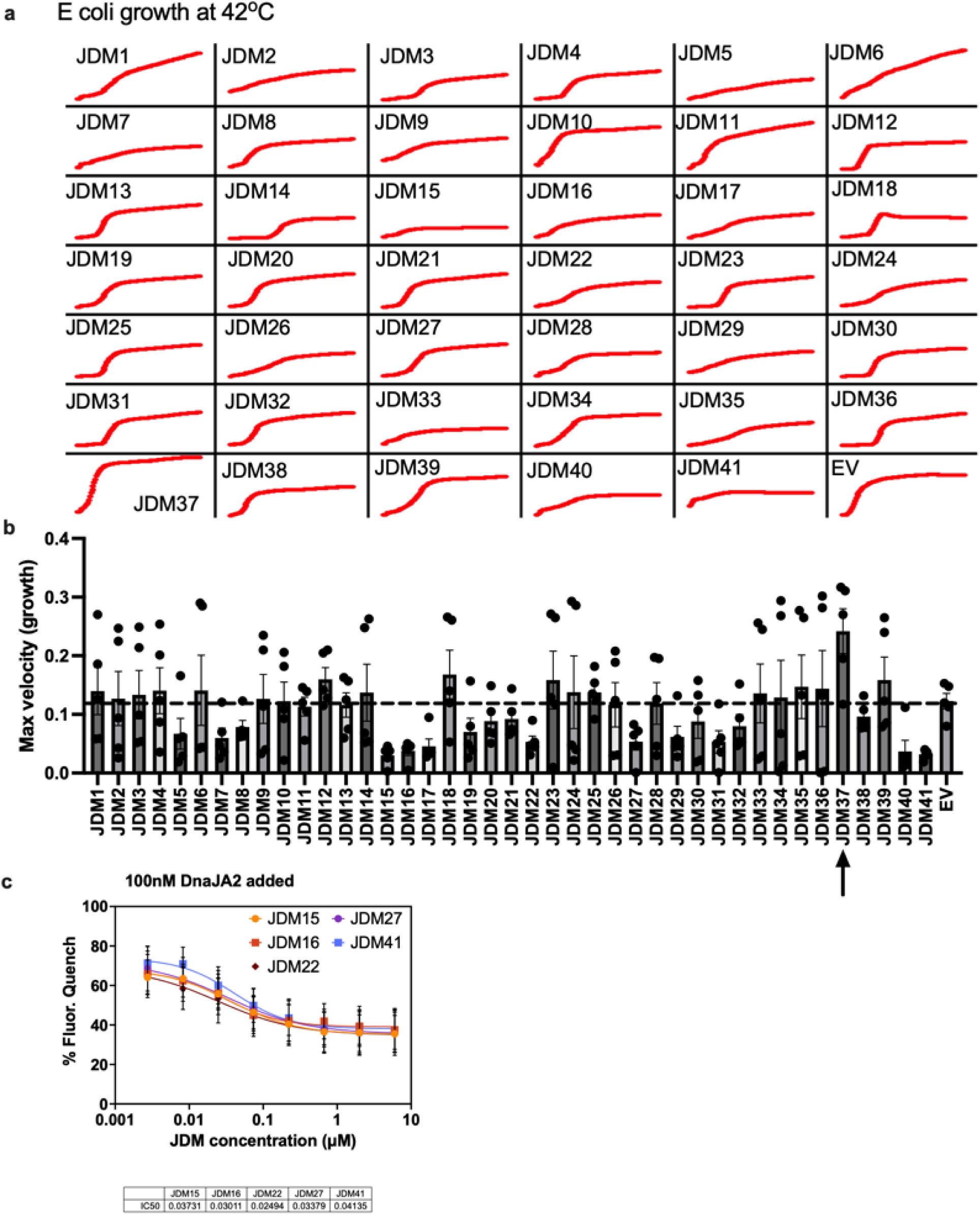
Measuring the effect JDMs have on *E. coli.* cell growth. **a-b**, The top 41 final JDM designs were transformed into *E. coli.*, induced for expression, grown at 42°C, and their cell growth was tracked. Shown is a representative cell growth curve (**a**) or the average maximum growth velocity (**b**) (n=5 experiments). Dotted line is for empty vector expression as control. (**c**) The JDMs with the most consistent and dramatic inhibition of *E. coli* growth at 42°C were tested for inhibition of DnaJA2 (100nM) activation of Hsc70. IC50s are shown on the bottom with the units of µM.

**Extended Data Figure 7:**
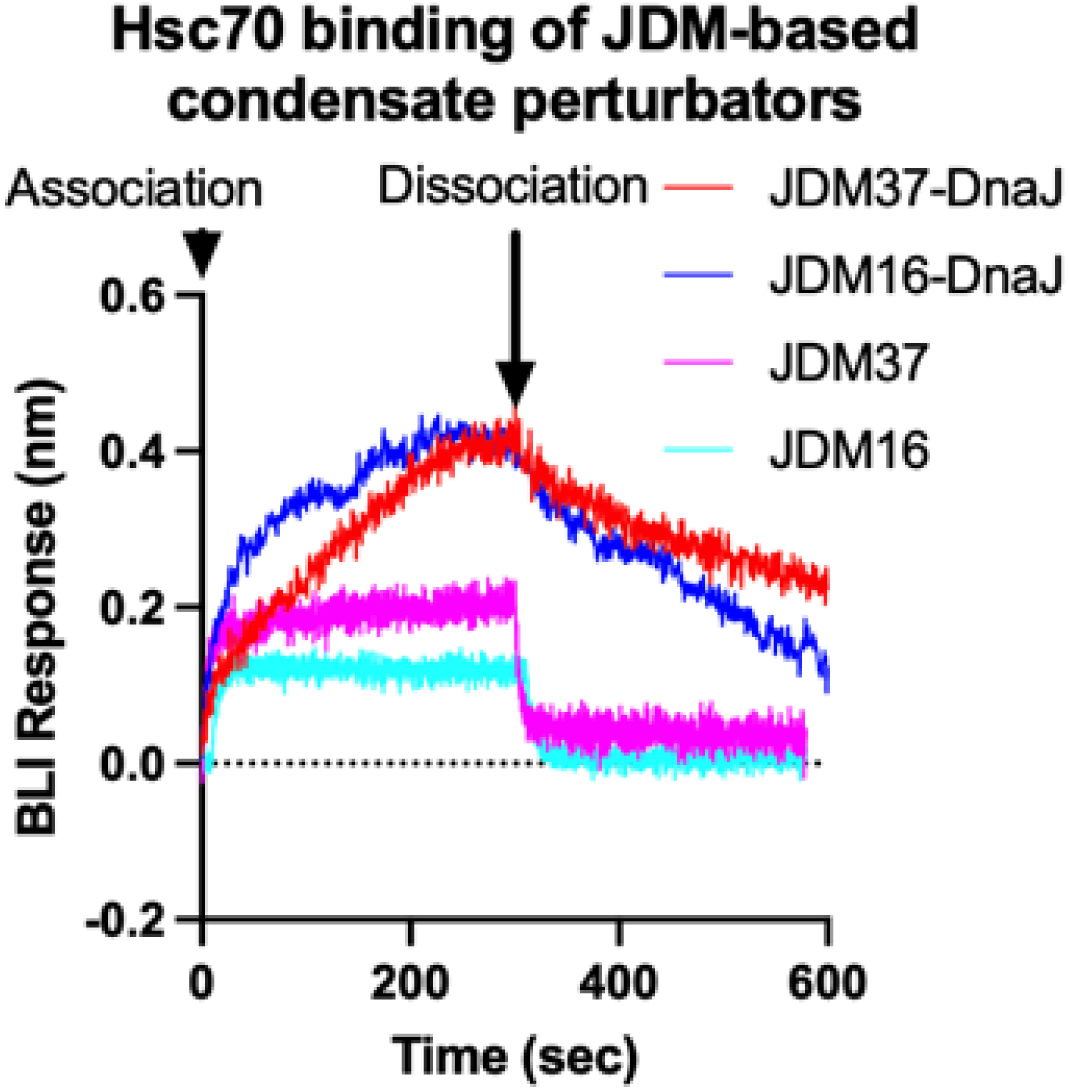
Measuring the binding of JDM-based condensate perturbators to Hsc70. Representative raw kinetic BLI data of JDM37-based or JDM16-based condensate perturbators binding to ATP-loaded Hsc70. JDM37 and JDM16 alone are displayed as comparisons. Concentration used for each protein displayed here is 729nM.

**Extended Data Figure 8:**
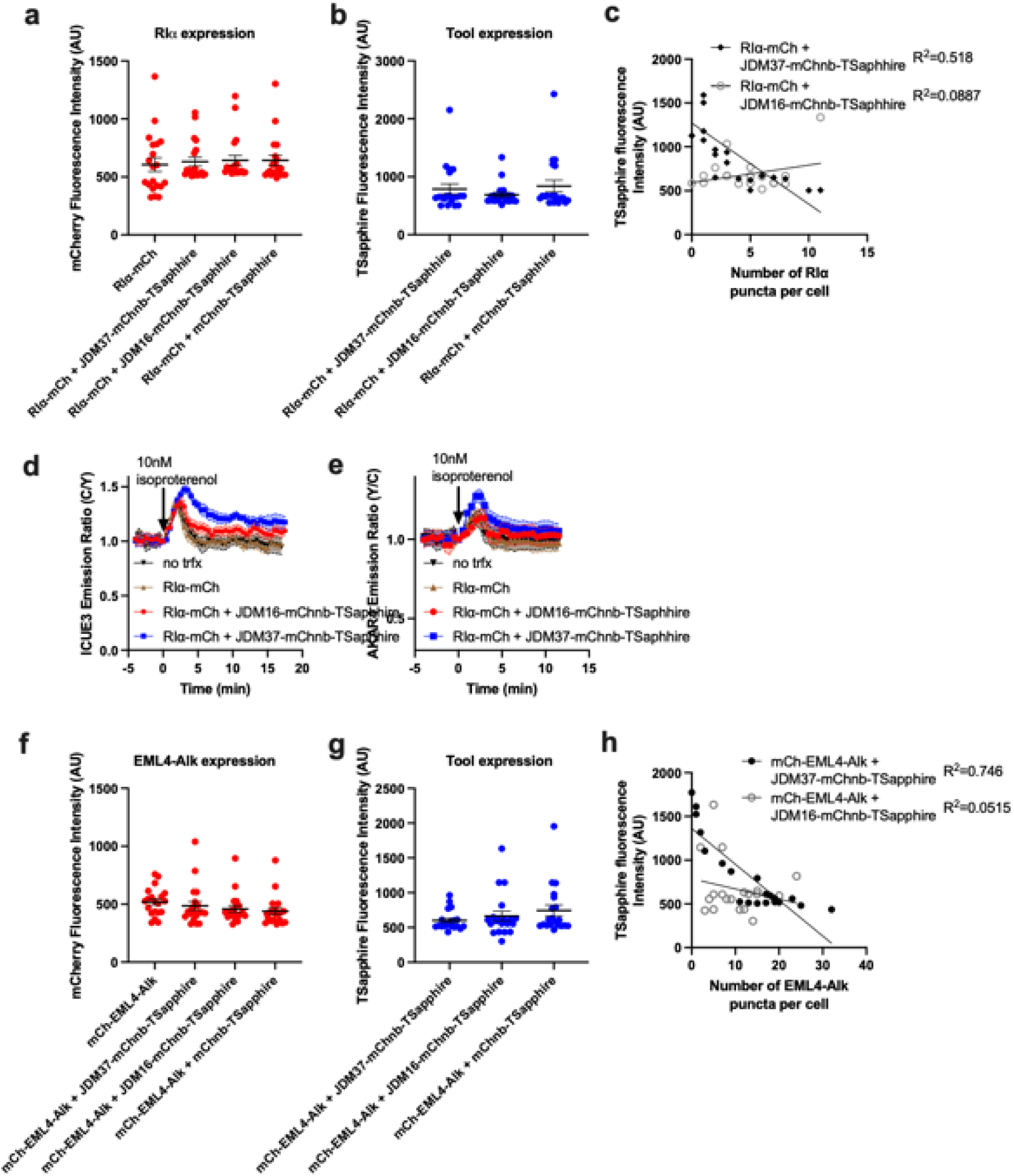
Fluorescence metrics of condensate perturbators expressed in mammalian cells. **a-b**, Average mCherry (**a**) or TSapphire (**b**) fluorescence intensity of different constructs tested in HEK293T cells. Each point represents a single cell (n=20 cells). **c**, Scatterplot comparing Tsapphire fluorescence intensity (condensate perturbator expression) to number of RIα puncta per cell. **d-e**, Time-course imaging of HEK293T cells expressing mCherry-tagged RIα, either cAMP sensor ICUE3 (**d**) or PKA sensor AKAR4 (**e**), and either JDM37-based or JDM16-based condensate perturbators. Also no transfection and mCherry tagged RIα were done as controls. In each condition, 10nM isoproterenol was added. Solid lines indicate representative average time with error bars representing standard error mean (SEM) (n=at least 15 cells per curve). **f-g**, Average mCherry (**f**) or TSapphire (**g**) fluorescence intensity of different constructs tested in Beas2B cells. Each point represents a single cell (n=20 cells). **h**, Scatterplot comparing Tsapphire fluorescence intensity (condensate perturbator expression) to number of EML4-Alk puncta per cell.

**Extended Data Figure 9:**
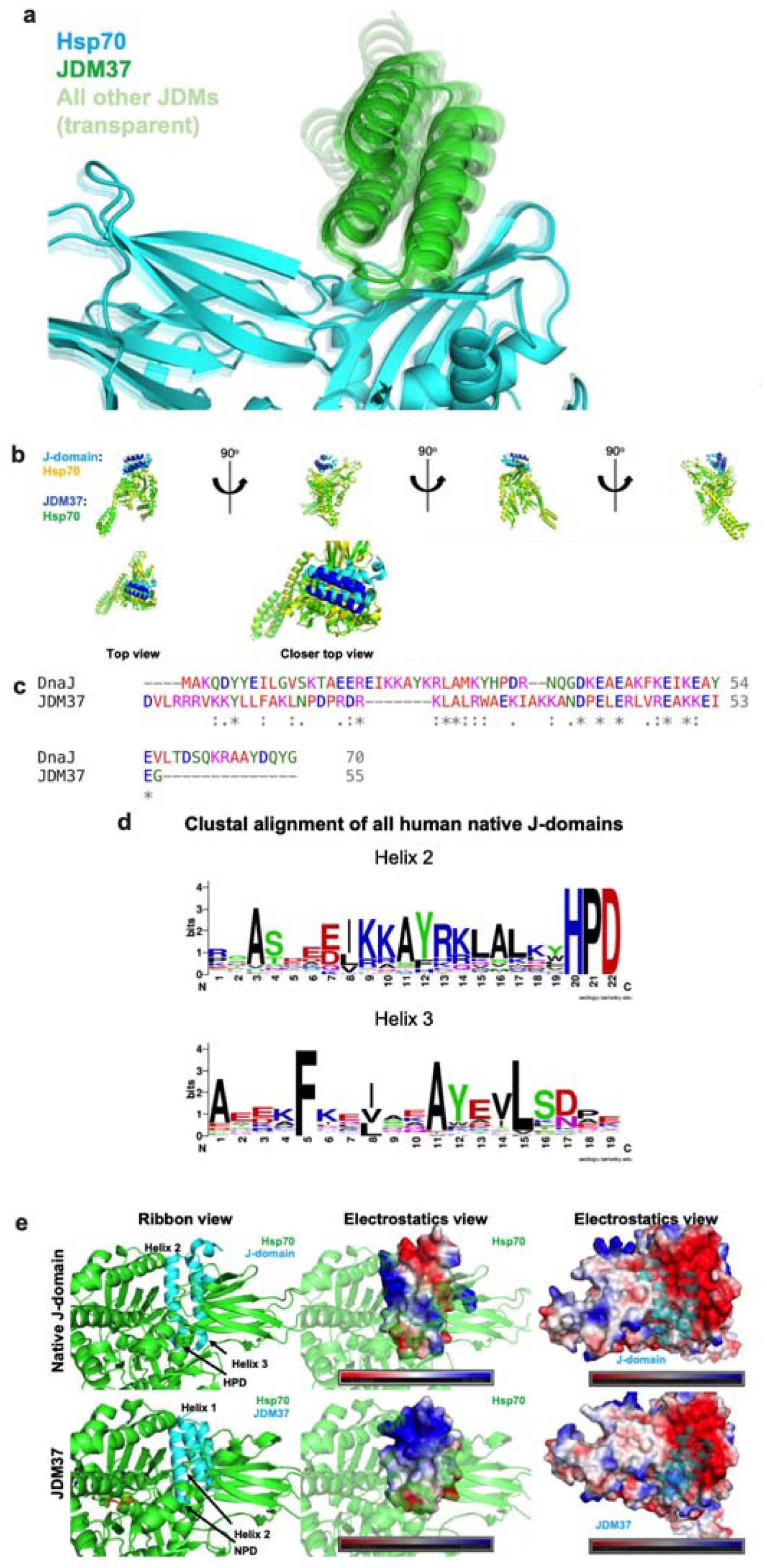
Exploring the sequence and structural determinants for activating Hsp70. **a**, AlphaFold2 predictions of the top 41 JDMs in complex with Hsp70. JDM37 is solid green while the remaining JDMs are transparent. **b**, Multiple viewpoints of either native J-domain:Hsp70 (cyan:yellow) (PDB: 5NRO) or JDM37:Hsp70 complex (blue:green). **c**, Sequence alignment of native J-domain from DnaJ versus JDM37. **d**, Clustal alignment of all human native J-domains. **e**, Various maps of either native J-domain:Hsp70 (top) or JDM37:Hsp70 (bottom) complex. Left: ribbon representation, Middle: electrostatic map of native J-domain or JDM37, Right: electrostatic map of Hsp70.

**Supplementary Note: Exploring the sequence and structural determinants for activating Hsp70**

While binding is not sufficient for Hsp70 activation (**Fig. 1b**), we explored if there are structural features unique to JDM37 to gather information about the determinants in activating Hsp70. Using AlphaFold2 structure predictions and crystal structures of DnaJ:DnaK complex, we explored whether structure can discern Hsp70 activators compared to inhibitors. However, there were no clear differences between JDM37 and the other JDMs or native J-domains in terms of Hsp70 interactions and conformation (**Extended Data Fig. 9a-b and Table S1**), although many of these complexes are only predictions. While there is very little sequence similarity between JDM37 and Hsp70 including the absence of the highly conserved HPD motif, JDM37 does have a very similar NPD motif (**Extended Data Fig. 9c-d**). Moreover, the electrostatics involved in Hsp70 binding (helices 2 for native J-domain and JDM37) are also largely conserved (**Extended Data Fig. 9e**). However, JDM37 and the other JDMs have high sequence similarity especially in their charged residue patches (**Extended Data Fig. 2**), thus the pattern of these electrostatic patches is also conserved for the other JDMs. Thus, it is still not obvious what differentiates JDM37 when looking at binding, structure, or sequence so far.

**Table S1: Rosetta metrics of JDM:DnaK predicted complexes**

Using AF2, each JDM was assessed for their predicted binding to DnaK (*E. coli* Hsp70). Afterwards, each JDM:DnaK complex was scored based on Rosetta and AF2 metrics.

## Notes

### Competing Interest Statement

The authors have declared no competing interest.

